# Sex differences along the Autism Continuum: A twin study of brain structure

**DOI:** 10.1101/330225

**Authors:** Élodie Cauvet, Annelies van’t Westeinde, Roberto Toro, Ralf Kuja-Halkola, Janina Neufeld, Katell Mevel, Sven Bölte

**Affiliations:** Center of Neurodevelopmental Disorders at Karolinska Institutet (KIND), Division of Neuropsychiatry, Department of Women’s and Children’s Health, Karolinska Institutet, Stockholm, Sweden; Center for Psychiatry Research, Stockholm County Council, Stockholm, Sweden; Human Genetics and Cognitive Functions, Institut Pasteur, Paris, France; CNRS URA 2182 “Genes, synapses and cognition”, Paris, France; Université Paris Diderot, Sorbonne Paris Cité, Human Genetics and Cognitive Functions, Paris, France; Department of Medical Epidemiology and Biostatistics, Karolinska Institutet, Stockholm, Sweden; Laboratory for the Psychology of Child Development and Education, CNRS Unit 8240, Paris-Descartes University and Caen University, Alliance for higher education and research Sorbonne Paris Cité (IDEX), Sorbonne, France; Child and Adolescent Psychiatry, Stockholm County Council, Sweden

**Author notes:** Corresponding Author: Élodie Cauvet, Child and Adolescent Psychiatry Research Center, KIND, Gävlegatan 22B, S-113 30 Stockholm, Sweden. Equal contribution (Élodie Cauvet, Annelies van’t Westeinde).

**Keywords:** Autism Spectrum Disorder, Autistic Traits, Biological Sex, Neuroanatomy, Twins

## Abstract

Females might possess protective mechanisms regarding autism spectrum disorder (ASD) and require a higher detrimental load, including structural brain alterations, before developing clinically relevant levels of autistic traits. This study examines sex differences in structural brain morphology in autism and autistic traits using a within-twin pair approach. Twin design inherently controls for shared confounders and enables the study of gene-independent neuroanatomical variation. N=148 twins (62 females) from 49 monozygotic and 25 dizygotic same-sex pairs were included. Participants were distributed along the whole continuum of autism including twin pairs discordant and concordant for clinical ASD. Regional brain volume, surface area and cortical thickness were computed. Within-twin pair increases in autistic traits were related to decreases in cortical volume and surface area of temporal and frontal regions specifically in female twin pairs, in particular regions involved in social communication, while only two regions were associated with autistic traits in males. The same pattern was detected in the monozygotic twin pairs only. Thus, non-shared environmental factors seem to impact female more than male cerebral architecture. Our results are in line with the hypothesis of a female protective effect in autism and highlights the need to study ASD in females separately from males.

---

Autism spectrum disorder (ASD) is a neurodevelopmental condition characterized by impairments in social interaction and communication alongside stereotypic, restricted, and repetitive behavior and interests, causing functional impairment in everyday life (American Psychiatric Association 2013; de Schipper et al. 2015) ASD constitutes the extreme end of a continuous distribution of autistic traits (Ronald et al. 2006; Robinson et al. 2016), and is associated with neuroanatomical alterations (Ecker et al. 2015). The precise nature of such alterations has remained obscured hitherto (Haar et al. 2014), partly due to the heterogeneous etiology and phenotypic expression of ASD (Katuwal et al. 2016). With an estimated heritability of 50-95% (Sandin et al. 2014; Colvert et al. 2015), the phenotypic variation of ASD is associated with a multitude of genetic and environmental factors (Ronald et al. 2006), likely resulting in a heterogeneous brain anatomy.

Twin designs are appropriate to account for heterogeneity when assessing ASD-related structural brain alterations. The twin co-twin control design enables to study associations between brain structure, autism and autistic traits, using the co-twin as the perfect control on age, gender, socio-economic background, other shared-environment and half (dizygotic) or all (monozygotic) genetic make-up. Further, restricting the analyses to monozygotic (MZ) twin pairs allows for the evaluation of non-shared environmental influences. More specifically, these analyses enable to study gene-independent neuroanatomical variation, thus pointing to the possible existence of environmental factors influencing pathological brain development. In the end, detecting such factors would provide possibilities for altering the developmental course of ASD. To date, only a handful of twin studies on ASD neuroanatomy have been conducted on small samples of twin pairs, reporting widespread differences across the brain (Mevel et al. 2014).

None of the previous twin-studies of ASD has addressed sex differences, and generally females have been neglected in neuroanatomical autism research (Via et al. 2011). The latter is unfortunate, since the skewed sex ratio in ASD, varying between 4:1 and 2:1 (Werling and Geschwind 2013; Baxter et al. 2015; Idring et al. 2015; Lai et al. 2015), points at the relevance of investigating differences in male and female brains. Sex-related differences might involve increased relative risk for males, and a protective effect in females (Werling and Geschwind 2013). Indeed, having a “male-like” neuroanatomical phenotype has recently shown to be associated with increased risk for ASD, while having a “female-like” neuroanatomical phenotype was associated with reduced risk for ASD (16). A protective effect of a female-like neuroanatomy might entail that more structural changes are required before reaching similar levels of autistic symptomatology in females, as hypothesized by the “multi-factorial sex/gender-differential liability model” (Werling and Geschwind 2013; Lai et al. 2015). Therefore, the inclusion of females exhibiting sub-threshold broader autistic phenotypes in research is crucial. In addition, the neuroanatomy related to ASD in females might differ from males in its location and nature. For example, decreased cortical thickness is observed in females with ASD, as opposed to increased cortical thickness in males with ASD, in the inferior and middle temporal lobes (Ecker et al. 2017). Therefore, both quantitative and qualitative neuroanatomical differences in association with autistic traits might be expected between males and females, as previous research indeed indicates (Bloss and Courchesne 2007; Craig et al. 2007; Schumann et al. 2010; Lai et al. 2013; Schaer et al. 2015; Retico et al. 2016). Using a within-pair design provides valuable insights into neuroanatomical sex differences by testing if autistic trait differences within female twin pairs are associated with more structural brain differences compared to male twin pairs. Further, by assessing autistic traits in twins, in addition to only clinical diagnosis of ASD, the ascertainment bias of possibly underdiagnosing females is avoided and females with sub-threshold clinical phenotypes are included. Such an approach is also consistent with the continuous distribution of autistic traits in the general population (von dem Hagen et al. 2011; Ecker et al. 2015) and in line with recent psychiatry research paradigms recommending dimensional constructs to study behaviors from typical to atypical (RDoC - Research Domain Criteria, NIH).

The current study sought to assess, for the first time, sex differences in structural morphology related to autistic trait severity in twins. We used a within pair design in a twin cohort of individuals with ASD, other neurodevelopmental conditions and typical development collected from the Roots of Autism and ADHD Twin Study Sweden (RATSS) (Bölte et al. 2014). An atlas-based parcellation approach was implemented using both volume- and surface-based morphometry to identify brain regions showing significant within-pair associations between autistic traits and brain morphometry estimates, including gray and white matter volumes, cortical thickness and surface area.

## Materials and Methods

### Participants

A sample recruited from RATSS was studied, including n=95 same-sex twin pairs for which complete MRI T1 scans were available. After excluding pairs with low image quality, n= 74 twin pairs (31 female twin pairs, mean age 16.2 yrs.; 43 male pairs, mean age 15.5 yrs.) were retained for the surface-based analyses. These included 49 monozygotic pairs and 25 dizygotic pairs. Zygosity was mostly based on DNA testing, apart from 2 female and 2 male pairs, for which zygosity was derived from a questionnaire. In RATSS, the twins are either discordant or concordant for ASD diagnosis or other neurodevelopmental conditions, or concordant for typical development. The sample studied here included 5 female- and 11 male ASD discordant pairs, 3 female- and 3 male ASD concordant pairs, and 23 female and 29 male pairs without ASD. Of the n=148 subjects, 28 received a clinical diagnosis for ASD (11 females, 17 males). The complete sample composition and demographics for the surface-based analyses are presented in Table 1 and Supplementary Table 5 with diagnostics details.

**Table 1:**
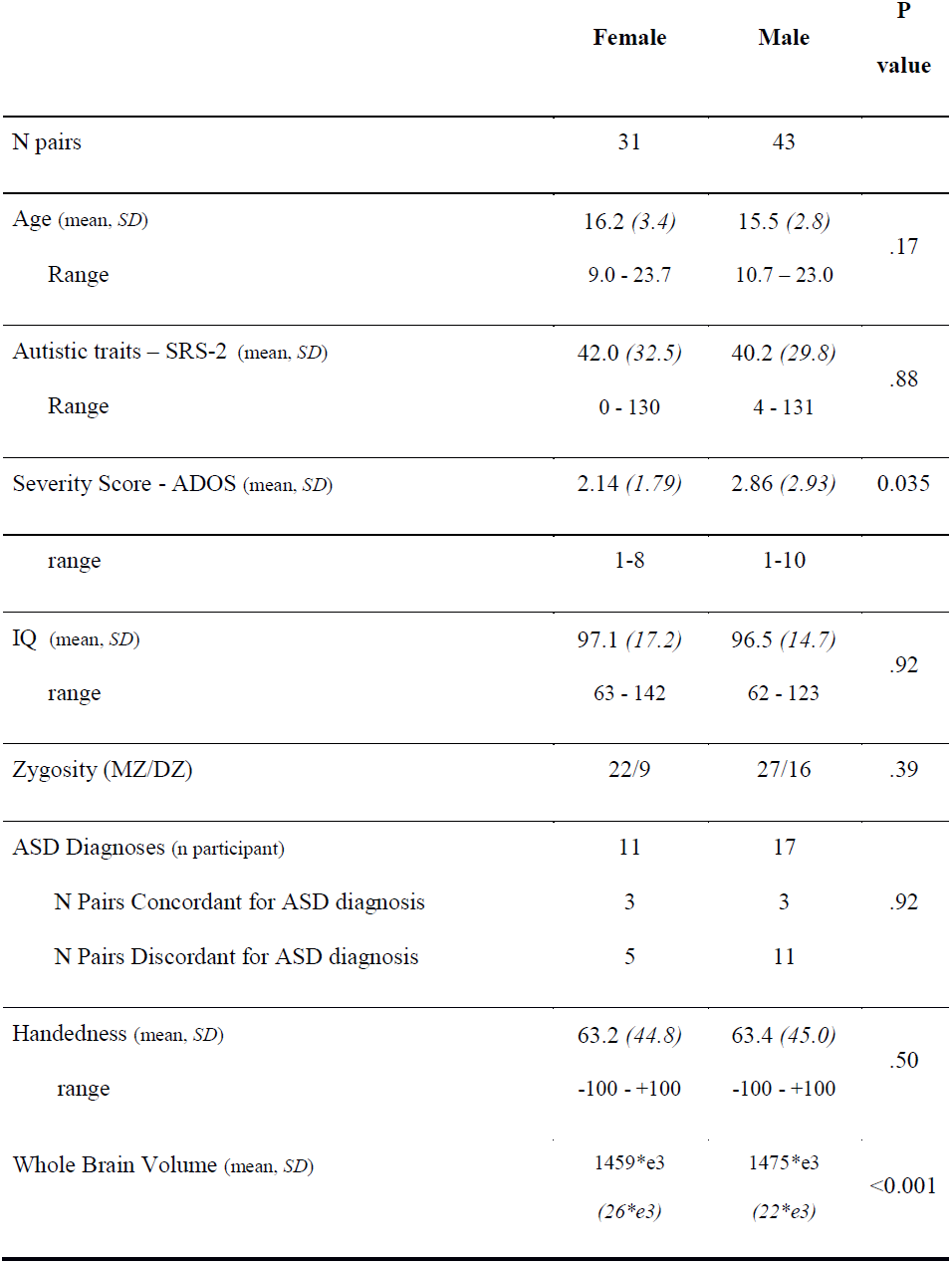
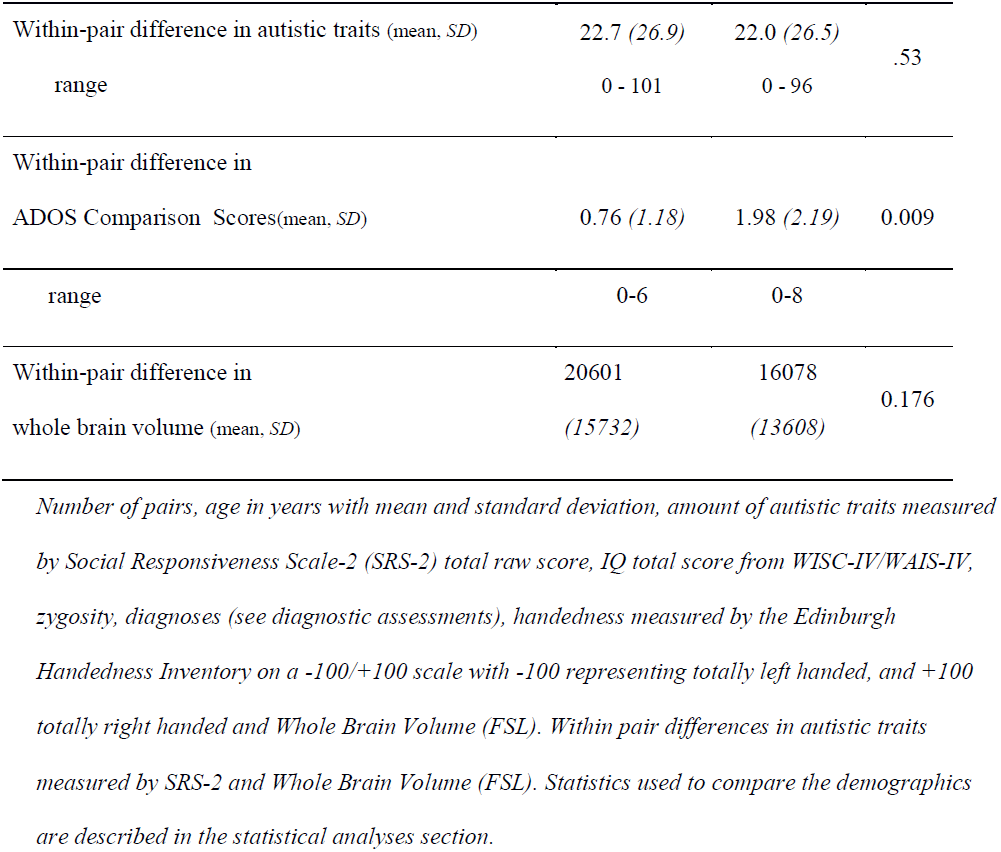
Sex-specific sample characteristics for surface-based morphometry analysis

### Procedures

Informed written consent was obtained from all participants and/or their legal guardians. The RATSS project is approved by the Regional Ethical Review Board.

#### Diagnostic and behavioral assessments

Twins were assessed according to a comprehensive psycho-diagnostic protocol (Bölte et al. 2014). Clinical consensus diagnosis was based on DSM-5 criteria (American Psychiatric Association 2013) by three experienced clinicians, supported by findings from the Autism Diagnostic Interview – Revised (Rutter et al. 2003), the Autism Diagnostic Observation Schedule-2 (Lord 2012), the Kiddie Schedule for Affective Disorders and Schizophrenia (Kaufman et al. 1997) or the Diagnostic Interview for ADHD in Adults (Kooij 2010). Full-scale IQ was assessed based on the Wechsler Intelligence Scales for Children or Adults, Fourth Editions (Wechsler 2003, 2008). Handedness was assessed using the Edinburgh Handedness Inventory (Oldfield 1971), which estimates laterality in everyday life on a scale from -100% (left handed) to +100% (right handed).

#### Autism trait and symptom severity measures

The Social Responsiveness Scale-2 (SRS-2) (Constantino 2012) assesses autistic-like behaviors and quantifies their severity during the past six-month, operationalizing social communication, social motivation, social awareness, social use of language, and rigid inflexible behaviors. It comprises 65 Likert-scaled items scored 0 to 3 generating a total score ranging from 0 to 195, with higher scores indicating more autistic traits. The SRS-2 has demonstrated good to excellent psychometric properties and superior accuracy to other measures of autistic traits (Bölte et al. 2011). Total raw scores from the parent-report SRS-2 standard child or adult versions were used to determine the extent of autistic traits, as recommended for research settings (Constantino 2012). A cut-off score of > 75 is recommended for clinically relevant expressions of autism. A Kolmogorov-Smirnov test revealed that total raw score distributions did not differ between the child and adult versions (D=.26, p=.20).

In addition to traits, we measured clinical autism symptom severity with ADOS comparison scores. The ADOS consists of different administration variants (modules), depending on developmental and language level, combining structured play and interview elements, and requiring about 30 to 60 minutes to conduct. The objective of the ADOS is to provide a context where social-communication and repetitive, restricted behaviors relevant to the diagnosis of ASD can be observed with high likelihood. ADOS comparison scores are ASD severity scores, comparable across modules, ranging from 1 to 10, with 1 indicating minimal or no symptoms, and 10 indicating severe symptoms of ASD.

### Structural MRI

#### Image Acquisition

T1-weighted images were acquired on a 3 Tesla MR750 GE scanner at the Karolinska Institutet MR center (Inversion Recovery Fast Spoiled Gradient Echo - IR-FSPGR, 3D-volume, 172 sagittal slices, 256×256, FOV 24, voxel size 1mm3, flip angle 12, TR/TE 8200/3.2, using a 32-channel coil array). T1-weighted acquisition was part of a 50-minute scanning protocol preceded by 5 to 7 minutes of MOK-scan training. During MOK-scan training, participants’ head movement was measured with a sensor on the forehead, and feedback was provided through a movie that stopped if the subjects moved too much.

#### Image processing

All images were processed in FSL 4.1 and Freesurfer 5.3. Total brain volume estimations were obtained in FSL, while regional cortical volumes, cortical thickness and surface area were obtained for the 148 parcels from the Destrieux Atlas(Destrieux et al. 2010) in Freesurfer.

#### Brain volume extraction (FSL)

The raw volumes were intensity normalized and the brain was extracted using AFNI’s 3dskullstrip. Skull stripped 3D images were segmented in 3 tissue compartments (Gray Matter-GM, White Matter-WM, Cerebral Spinal Fluid-CSF) using FAST (FMRIB’s Automated Segmentation Tool), which also corrects for spatial intensity variation. Segmented images were warped to MNI space using non-linear registration FNIRT from FMRIB’s Software Library. Gray and white matter compartment volumes where then summed for each individual to get the total brain volume. Regional brain volume analyses in FSL were also conducted, without leading to significant results (see supplementary material).

#### Surface-based: cortical volumetry, cortical thickness and surface area (Freesurfer)

Images rated as good quality after FSL processing were also processed in Freesurfer (http://surfer.nmr.mgh.harvard.edu/). The standard well validated pipeline was run on the raw T1-weighted images (Dale et al. 1999; Fischl et al. 1999) to derive cortical thickness (mm), grey matter volume (mm^3^) and surface areas (mm^2^) for each region from the Destrieux Atlas (Destrieux et al. 2010). A quality control procedure was done in which 2 researchers visually inspected the images to check that the pial- and white matter surface line accurately followed the grey/white and grey/CSF borders throughout the brain. After this check, of the 158 processed individuals, 148 were retained for the surface-based analyses. Fronto-orbital regions and temporal poles have been excluded from the analyses because of segmentation errors. See supplementary material for details.

### Statistical Analysis

#### Sex differences in demographics and total brain volume

To rule out confounders as the source of potential neuroanatomical differences, we tested sex-differences using χ^2^ tests for categorical variables (zygosity, diagnosis) and Kruskal-Wallis tests for continuous variables (age, SRS-2, IQ, handedness scores, whole brain volumes [FSL]). Within twin-pair differences in autistic traits and clinical severity were computed for each twin pair. Kruskal-Wallis tests were run to assess if sex-differences existed in these within-pair difference scores, i.e. if females within a pair are more different than males within a pair. See figure 1, A and B for SRS-2 distributions.

**Caption Figure 1:**
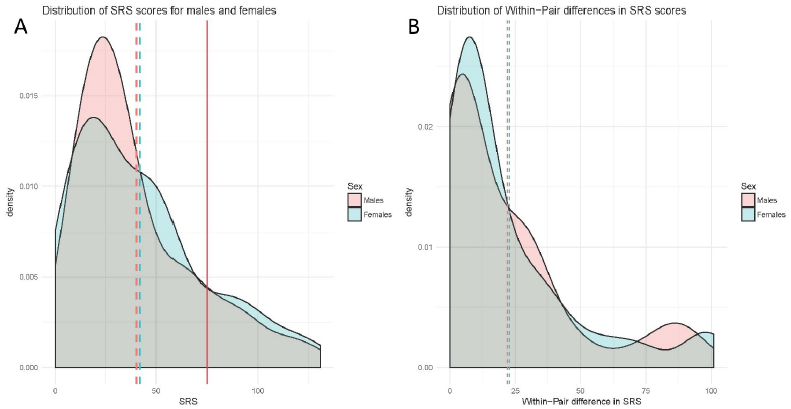
Population distribution of autistic traits (SRS-2 raw scores) as function of sex. B. Distribution of within pair differences in autistic traits. In red: Males, in blue: Females. Vertical dotted lines represent the mean for each group (males and females). Vertical full line represents the high cut-off of 75.

#### Within-pair analyses: neuroanatomy in relation to autistic traits

A within-pair (co-twin comparison) approach was used to estimate the association between autistic traits (exposure) and neuroanatomical measures (outcome) while controlling for unmeasured confounding factors shared within twin pairs (e.g. genetic factors, demographics etc.). Within-twin pair associations were estimated using a conditional linear regression model within the generalized estimating equations framework, using the drgee package from R (Zetterqvist and Sjölander 2015). Herein, the difference in the exposure variable within a pair is correlated to the difference in the outcome variable within the same pair (Figure 2A). Within-pair associations between autistic trait severity and brain structure were calculated for volumes of 96 regional gyri of gray and white matter from Harvard-Oxford in FSL as well as for cortical volume, surface area and thickness of 148 regions from the Destrieux Atlas in Freesurfer. Full-scale IQ and handedness were included as covariates in all analyses (see supplementary material). Whole brain volume, calculated from volume-based analysis (FSL), was added as a covariate only to the volume and surface area analyses, since it significantly impacts volume and surface area, but not cortical thickness (Toro et al. 2008).

**Caption Figure 2:**
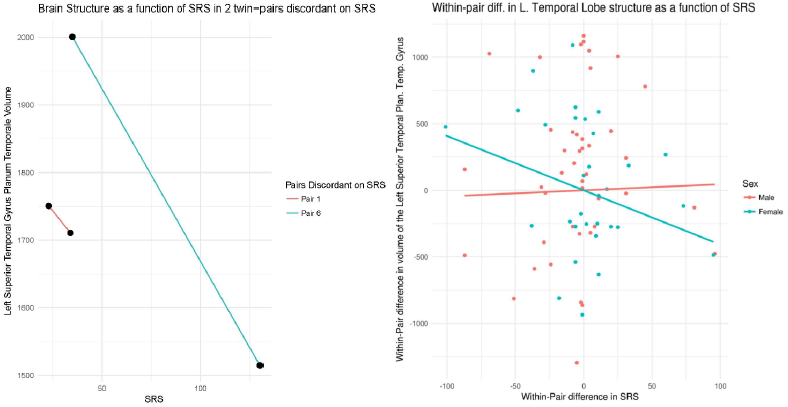
A. Example in two qualitatively discordant female pairs of differences in cortical volume in function of their autistic traits (SRS-2 total score) in the left superior temporal gyrus Planum Temporale, illustrating the within pair negative association between regional brain volume and autistic traits. B. Illustration of within pair association between cortical volume and autistic traits (SRS-2 total score) in females (blue) and males (red) for the unadjusted model (i.e. without full scale IQ, handedness and total brain volume). The model reported in this paper included all covariates. Each dot represents a pair. The line corresponds to the estimate from the regression of within pair differences in neuroanatomical in function of differences in autistic traits.

#### Step-wise group analyses

Whole group within-pair associations between brain structure and autistic trait severity were first calculated without taking sex into account. Second, sex was added as a factor in the model so that separate estimates for males and females are retrieved. Third, a χ^2^ test assessed if the number of regions that were statistically significantly associated with autistic traits was different between males and females (quantitative sex difference). Finally, for all the regions that were significantly associated with autistic traits in either males or females, we calculated Wald-tests to investigate if the estimates of the associations were different between the sexes (qualitative sex difference).

#### Post-hoc analyses

Additional within-pair analyses were run on the regions significantly associated with autistic traits. First, to test the influence of non-shared environmental factors on the associations, the sample was restricted to MZ twin pairs, removing genetic differences. Second, to assess if the association between autistic traits and brain structure was not only driven by clinical and high-trait cases, but equally valid for the middle and lower end of the autism continuum and subclinical broader phenotypes, we re-ran the analysis on a subset of subjects excluding pairs in which at least one displayed high-autistic traits (>75) or had an ASD diagnosis. Further, we explored the association of brain structure with clinical severity using ADOS scores. Finally, to study the interaction between age and autistic traits on brain structure, we conducted a standard linear regression model across twin pairs, with brain structure as outcome, and autistic traits, brain volume, IQ, age and sex as predicting variables. Previous research has shown a linear age effect in our age-group, therefore, no quadratic effects were assessed (Shaw et al. 2008). Cluster robust standard errors (“sandwich”) was used to account for heteroscedasticity, non-normal distribution, and pair clustering of the data.

All statistical analyses were performed in R (www.r-project.org). The p-values for the main analyses were FDR corrected for Type I errors and significance threshold set to p<.05. The FDR correction was based on 148 regions of interest, and done separately for each estimate (volume, surface area and thickness). The post-hoc analyses on the significant regions from the main analyses are uncorrected, with a significance threshold of p <.05.

## Results

### Sex differences in demographics and total brain volume

No significant differences between males and females were observed for age, autistic traits (SRS-2), IQ, and handedness scores (Table 1). Females had smaller whole brain volumes compared to males (χ^2^(1) =17.59, p<0.001), and lower autistic symptom severity scores (ADOS) (χ^2^(1) =4.44, p=0.035). Further, there were no sex differences on within-pair difference of most these variables, including autistic traits, i.e. female twins were not significantly more different from their co-twins on parent-rated autistic trait severity, compared to males. However, females showed a smaller within-pair difference on autism symptom severity on the ADOS compared to males (χ^2^(1) =6.74, p=0.009).

### Surface-based regional morphometry: cortical volume, cortical thickness and surface area

Table 2 reports significant within-pair associations between autistic traits and surface based-morphometry estimates including cortical volume, surface area and thickness. The significant within-pair associations, with autistic traits as a predictor of brain structure, are reported both for the sexes separately and for the whole sample. The whole-sample analyses yielded no significant result. However, when splitting the sample in males and females, several within-pair associations between autistic traits and brain structure were observed (Figure 3).

**Table 2:**
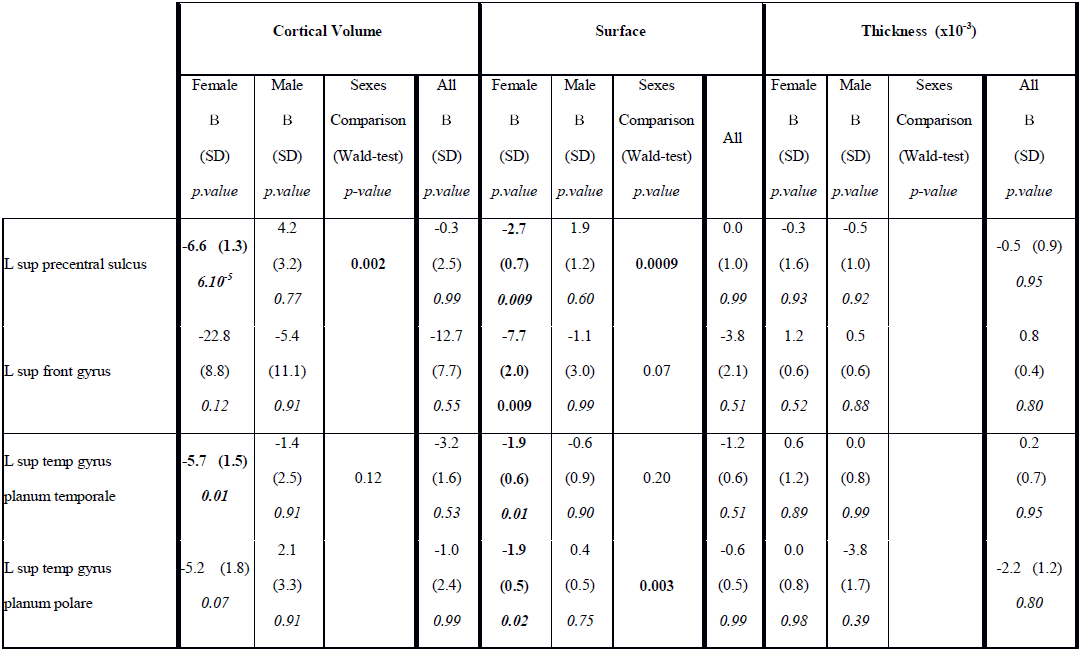
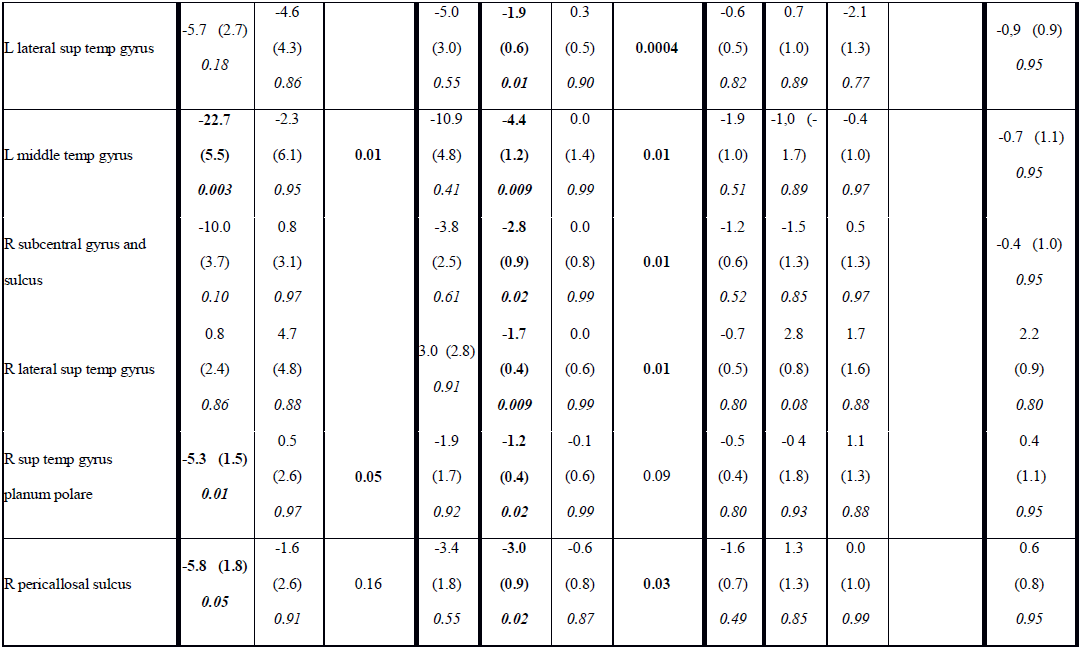
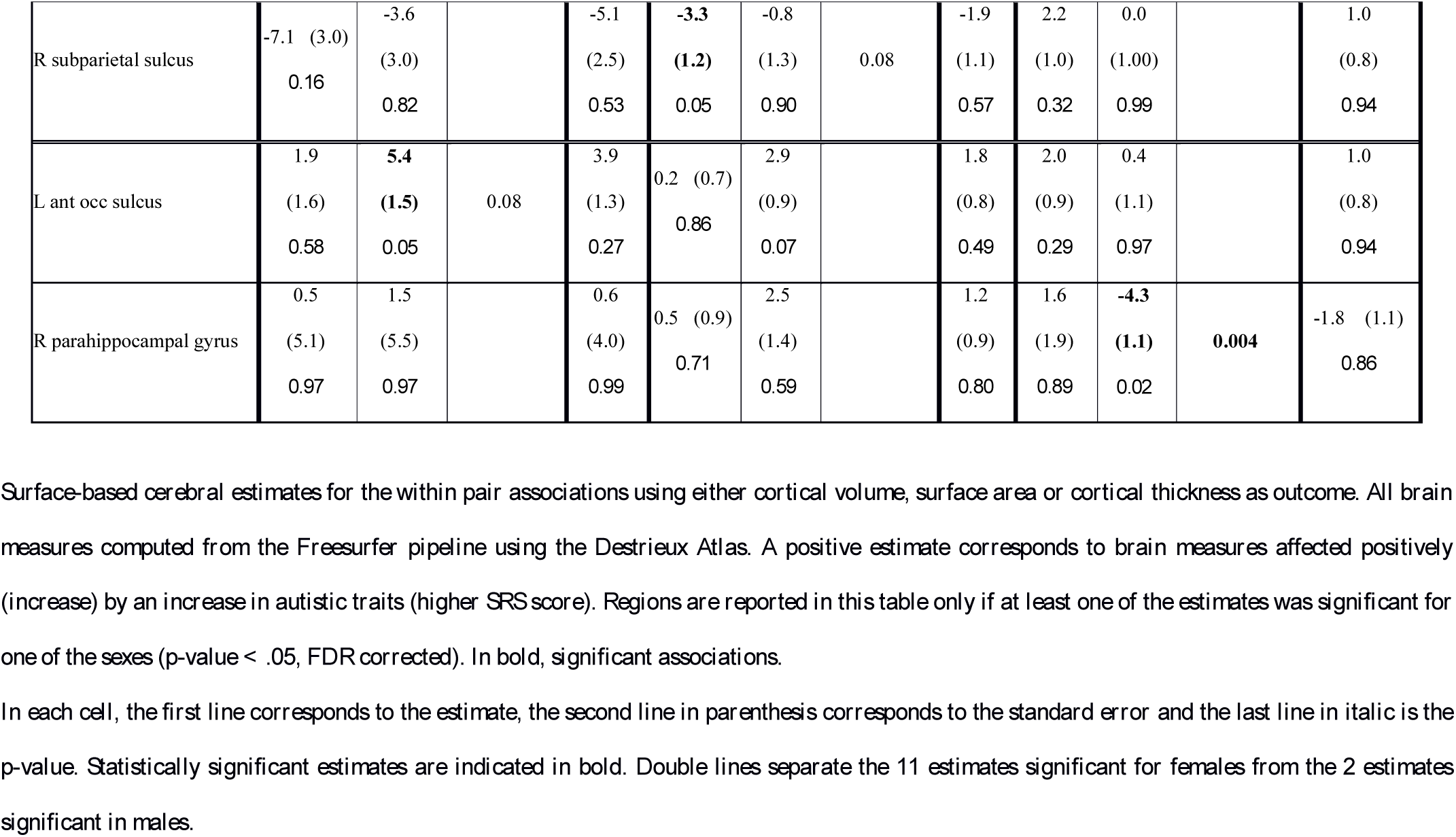
Within-pair associations between autistic traits and surface-based cerebral estimates (n=148, 62 females and 86 males)

**Caption Figure 3:**
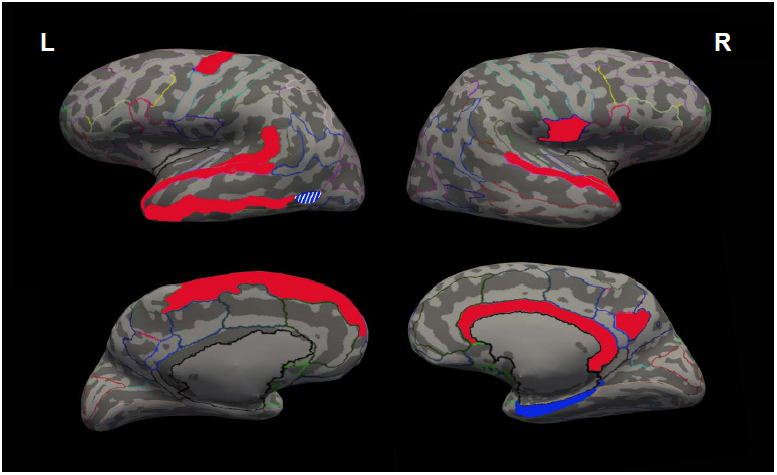
Illustration of surface based morphometric results displayed on inflated brain. In red: regions with significant association between autistic traits and brain estimates in females; in blue: regions for males; plain colors: negative correlations; stripped colors: positive correlations (only in left anterior occipital sulcus for males). L: left hemisphere; R: right hemisphere. Upper part: lateral side, lower part: medial side.

In females, a within-pair increase in autistic traits was associated with a within-pair reduction of cortical volume and surface area of temporal and frontal gyri and sulci. These included the left superior temporal gyrus planum temporale, middle temporal gyrus, lateral superior temporal gyrus and superior precentral sulcus, as well as the right superior temporal gyrus planum polare and pericallosal sulcus (Table 2). Further, in females, a within-pair increase in autistic traits was associated with a decrease in surface area in the left superior frontal and right lateral superior temporal gyri, in addition to a trend for increase in thickness of the latter region, resulting in no significant change in volume. Thus, within the female pairs, the twins with the higher autistic traits had reduced volume and surface area compared to their co-twins in most of these regions. An example of the within-pair association between increased traits and reduced volume of the left superior temporal gyrus planum temporale in 2 female twin pairs is shown in figure 2A. Figure 2B displays the same association across the male and female cohorts. In males, on the other hand, a different pattern of associations was observed. A within-pair increase in autistic traits in males was related to within-pair increases in volume of the left anterior occipital sulcus and decreased thickness of the right para-hippocampal part of the medial occipito-temporal gyrus.

No brain regions were linearly associated with age nor with the interaction between age and autistic traits across the whole sample (Supplementary Table 4).

### Quantitative and qualitative sex differences: Sex specific regional alterations

The number of brain regions associated with within-pair differences in autistic traits was significantly smaller in males compared to females (11 in females and the two in males: Wald χ^2^ (1) =9.6, p =.002), suggesting quantitative difference. Further, Wald-test on the 13 regions related to autistic traits revealed that the estimate of the association between autistic traits and brain structure was significantly different between males and females for most regions, suggesting qualitative sex differences in cortical volume and surface area of the left precentral sulcus and left middle temporal gyrus; surface area of the left lateral superior temporal gyrus, right lateral superior temporal gyrus and right pericallosal area; and thickness in the right parahippocampal gyrus (Table 2).

### Within-pair associations in monozygotic twins: the influence of non-shared environment (n=98, 44 females and 54 males)

In monozygotic twins, the within-pair associations between a reduction in volume and/or surface area and autistic traits were significant in all but four regions in females (Supplementary Table 1), suggesting that their neuroanatomy is influenced by non-shared environmental factors. On the other hand, the loss of an association with autistic traits in the other regions, namely the left superior temporal gyri temporale and polare, the left superior precentral sulcus and right subparietal sulcus suggests their sensitivity to genetic factors.

### Within-pair associations in the sub-threshold group (SRS < 75 & no ASD-diagnosis, n=94, 40 females and 54 males)

To assess the effect of ASD diagnosis and high autistic traits on the previous association in the 13 significant regions, post-hoc analyses in the twin sample containing only low trait pairs revealed that within-pair increases in autistic traits were no longer significantly associated with neuroanatomical changes in females, while in males a different pattern of association was observed. An increase of traits within male pairs was associated with a bilateral surface area increase in the lateral superior temporal sulci (Supplementary Table 2).

### Within-pair associations between ADOS-severity and brain structure (n=138, 58 females and 80 males)

Similar associations between severity and brain structure were observed for autism symptom severity (ADOS) compared to parent-rated autistic traits (SRS-2), in both females and males (Supplementary Table 3), at the exception of two regions in the left temporal lobe, and the right pericallosal and subparietal sulci, which were no longer associated with brain structure in the females (see supplementary results).

## Discussion

This study is the first to assess sex differences in structural brain alterations related to autistic trait as well as autism symptom severity, while adjusting for genetic and environmental factors shared in twin pairs. Increases in autistic traits within twin pairs were significantly associated to decreases in cortical volume and surface area of temporal and frontal lobe regions in female, but not male twin pairs. In males, surface area was affected by autistic traits only in occipital and parahippocampal regions. Importantly, male and female twin pairs differed neither in their overall level of autistic traits, nor in their within-pair discordance for autistic traits. The observations therefore indicate that for a similar increase in autistic traits and clinical severity, females presented with both distinct and more structural brain alterations compared to males.

### Quantitative and qualitative structural brain differences

Our results endorse the observation from a recent large cohort study, based on singleton subjects from the ABIDE consortium, showing that few structural brain alterations are related to ASD in males (Colvert et al. 2015). Here, we show that when females are assessed separately, more structural alterations related to autistic traits are found. We demonstrate that female twins with more autistic traits and higher clinical severity differ more in their structural brain anatomy from their less affected co-twins as compared to male twin pairs, corresponding to previous observations of a greater extend of structural brain alterations in autistic females compared to controls (Bloss and Courchesne 2007; Schumann et al. 2010; Ecker et al. 2017). Importantly, since we assessed traits as opposed to only diagnosis, our findings relate to severity across the autism continuum, thereby reducing potential ascertainment biases. Although our design did not directly test a female protective effect, a more affected female brain supports theories of increased liability, i.e. a higher neuroanatomical load being required in females for reaching similar levels of autism severity (Lai et al. 2013; Werling and Geschwind 2013). However, since we did not observe the same associations in a subset without ASD cases and high-trait twins, while such associations were present for clinical symptoms, brain alterations in females might be observed only when clinical symptoms reach a certain threshold. Therefore, the anatomical alterations are rather associated with clinical ASD, as opposed to traits across the continuum. However, it must be noted that this is an exploratory study and the absence of results in the sub-threshold sample might reflect a lack of power triggered by smaller within pair differences in this sample. Thus, confirmatory analyses using population based samples are required to replicate the observations especially within lower end of the autistic continuum.

Further, the structural alterations observed in females appeared in different brain regions as compared to males, in particular reduced volume of temporal and frontal cortices, which has been observed in adults (Craig et al. 2007; Ecker et al. 2017), as well as young children in a reversed pattern (Retico et al. 2016). Indeed, little overlap in brain structure alterations associated with ASD might exist between sexes (Lai et al. 2013). Because of suspected differences in behavioral autistic phenotype between males and females (e.g. social communication) (Van Wijngaarden-Cremers et al. 2014), sex-specific structural brain alterations might be specific for certain symptom domains.

### Genetics vs. environment

Our observations suggest that both genetic and non-shared environmental factors are involved in the association between brain structure and autistic traits in females. Indeed, the regions associated with autistic traits in the monozygotic subsample points toward their specific sensitivity to non-shared environmental factors. However, future research measuring the actual load and type of non-shared environmental factors would enable to conclude whether females are more sensitive to environmental factors, or actually experience a higher non-shared environmental load impacting on brain structure. Determining which environmental factors underlie these gene-independent neuroanatomical variations might lead to target for reducing pathological developmental process in females that are vulnerable for developing ASD. In contrast, our exploratory post-hoc analyses suggest that, for regions in which the association was no longer significant in monozygotic twins, the relationship between structure and autistic traits might be influenced by genetic factors. Indeed, the precentral sulcus and left superior temporal planum temporale gyrus are among the first regions to develop in the embryo and under strong genetic control (Lohmann et al. 2008). However, future research with higher statistical power should seek to replicate these observations, since the limited power in this smaller sub-group might explain the reduction of significant associations.

Our sample consisted of subjects with a rather wide age range of 8 to 24 years. Although age is implicitly controlled for in a within-pair design, we cannot exclude the possibility that developmental effects modulate the observed associations, in particular given the slightly older age of the females with an ASD diagnosis in our sample. However, post-hoc analyses using a general linear model revealed no interactions between age and autistic traits on brain structure. Finally, some ASD relevant regions, including orbitofrontal regions (Craig et al. 2007; Schaer et al. 2015), had to be excluded from analyses due to insufficient segmentation quality.

In conclusion, using a well-controlled within twin-pair dimensional design, the current study identified more and different structural brain alterations related to autistic traits and clinical autism severity in females compared to males, partly being associated with non-shared environmental factors. Future studies should address sex differences more specifically in relation to genetic and environmental factors. Importantly, our findings highlight the need of assessing the etiology and expression of ASD in women separately from men.

## Acknowledgments

We would like to thank warmly our twin participants as well as their parents for participating in the RATSS project. We are also very grateful for the work and contributions from our colleagues Kerstin Andersson, Anna Lange-Nilsson, Steve Berggren, Torkel Carlsson, Christina Coco, Andreas Fällman, Martin Hammar, Johanna Ingvarsson, Johan Isaksson, Elzbieta Kostrzewa, Therese Lindström, Anna Lövgren, Lynnea Myers, Soheil Mahdi, Lina Poltrago, Anna Råde, Anna Lia Sacerdoti, Kristiina Tammimies, Elin Vahlgren, Charlotte Willfors, and Eric Zander.

The study was funded by the Swedish Research Council, Vinnova, Formas, FORTE, the Swedish Brain foundation (Hjärnfonden), Stockholm Brain Institute, Autism and Asperger Association Stockholm, Queen Silvia Jubilee Fund, Solstickan Foundation, PRIMA Child and Adult Psychiatry, the Pediatric Research Foundation at Astrid Lindgren Children’s Hospital, Sällskapet Barnavård, Jerring Foundation, the Swedish Order of Freemasons, Kempe-Carlgrenska Foundation, Sunnderdahls Handikappsfond, the Innovative Medicines Initiative Joint Undertaking (grant agreement number 115300), which comprises financial contribution from the European Union’s Seventh Framework Programme (FP7 /2007 – 2013) and in-kind contributions from companies belonging to the European Federation of Pharmaceutical Industries and Associations (EU-AIMS).

## References

American Psychiatric Association. 2013. Diagnostic and statistical manual of mental disorders (DSM-5^®^). American Psychiatric Pub.

Baxter AJ, Brugha TS, Erskine HE, Scheurer RW, Vos T, Scott JG. 2015. The epidemiology and global burden of autism spectrum disorders. Psychol Med. 45:601–613.

Bloss CS, Courchesne E. 2007. MRI Neuroanatomy in Young Girls With Autism. J Am Acad Child Adolesc Psychiatry. 46:515–523.

Bölte S, Westerwald E, Holtmann M, Freitag C, Poustka F. 2011. Autistic Traits and Autism Spectrum Disorders: The Clinical Validity of Two Measures Presuming a Continuum of Social Communication Skills. J Autism Dev Disord. 41:66–72.

Bölte S, Willfors C, Berggren S, Norberg J, Poltrago L, Mevel K, Coco C, Fransson P, Borg J, Sitnikov R, Toro R, Tammimies K, Anderlid B-M, Nordgren A, Falk A, Meyer U, Kere J, Landén M, Dalman C, Ronald A, Anckarsäter H, Lichtenstein P. 2014. The Roots of Autism and ADHD Twin Study in Sweden (RATSS). Twin Res Hum Genet Off J Int Soc Twin Stud. 17:164–176.

Colvert E, Tick B, McEwen F, Stewart C, Curran SR, Woodhouse E, Gillan N, Hallett V, Lietz S, Garnett T, Ronald A, Plomin R, Rijsdijk F, Happé F, Bolton P. 2015. Heritability of Autism Spectrum Disorder in a UK Population-Based Twin Sample. JAMA Psychiatry. 72:415–423.

Constantino JN. 2012. (SRS™-2) Social Responsiveness Scale™, Second Edition | WPS. Los Angeles: Western Psychological Services.

Craig MC, Zaman SH, Daly EM, Cutter WJ, Robertson DM, Hallahan B, Toal F, Reed S, Ambikapathy A, Brammer M, others. 2007. Women with autistic-spectrum disorder: magnetic resonance imaging study of brain anatomy. Br J Psychiatry. 191:224–228.

Dale AM, Fischl B, Sereno MI. 1999. Cortical surface-based analysis. I. Segmentation and surface reconstruction. NeuroImage. 9:179–194.

de Schipper E, Lundequist A, Coghill D, de Vries PJ, Granlund M, Holtmann M, Jonsson U, Karande S, Robison JE, Shulman C, others. 2015. Ability and disability in autism spectrum disorder: A systematic literature review employing the international classification of functioning, disability and health-children and youth version. Autism Res. 8:782–794.

Destrieux C, Fischl B, Dale A, Halgren E. 2010. Automatic parcellation of human cortical gyri and sulci using standard anatomical nomenclature. NeuroImage. 53:1–15.

Ecker C, Andrews DS, Gudbrandsen CM, Marquand AF, Ginestet CE, Daly EM, Murphy CM, Lai M-C, Lombardo MV, Ruigrok ANV, Bullmore ET, Suckling J, Williams SCR, Baron-Cohen S, Craig MC, Murphy DGM, Consortium MRCAIMS (Mrc A. 2017. Association Between the Probability of Autism Spectrum Disorder and Normative Sex-Related Phenotypic Diversity in Brain Structure. JAMA Psychiatry. 74:329–338.

Ecker C, Bookheimer SY, Murphy DGM. 2015. Neuroimaging in autism spectrum disorder: brain structure and function across the lifespan. Lancet Neurol.

Fischl B, Sereno MI, Dale AM. 1999. Cortical surface-based analysis. II: Inflation, flattening, and a surface-based coordinate system. NeuroImage. 9:195–207.

Haar S, Berman S, Behrmann M, Dinstein I. 2014. Anatomical Abnormalities in Autism? Cereb Cortex.

Idring S, Lundberg M, Sturm H, Dalman C, Gumpert C, Rai D, Lee BK, Magnusson C. 2015. Changes in prevalence of autism spectrum disorders in 2001-2011: findings from the Stockholm youth cohort. J Autism Dev Disord. 45:1766–1773.

Katuwal GJ, Baum SA, Cahill ND, Michael AM. 2016. Divide and Conquer: Sub-Grouping of ASD Improves ASD Detection Based on Brain Morphometry. PLoS One. 11:e0153331.

Kaufman J, Birmaher B, Brent D, Rao U, Flynn C, Moreci P, Williamson D, Ryan N. 1997. Schedule for Affective Disorders and Schizophrenia for School-Age Children-Present and Lifetime Version (K-SADS-PL): initial reliability and validity data. J Am Acad Child Adolesc Psychiatry. 36:980–988.

Kooij JJ. 2010. Diagnostic interview for ADHD in adults 2.0 (DIVA 2.0). Pearson Assess Inf BV Amst. ed. Adult ADHD Diagn Assess Treat.

Lai M-C, Lombardo MV, Auyeung B, Chakrabarti B, Baron-Cohen S. 2015. Sex/gender differences and autism: setting the scene for future research. J Am Acad Child Adolesc Psychiatry. 54:11–24.

Lai M-C, Lombardo MV, Suckling J, Ruigrok ANV, Chakrabarti B, Ecker C, Deoni SCL, Craig MC, Murphy DGM, Bullmore ET, Consortium MRCA, Baron-Cohen S. 2013. Biological sex affects the neurobiology of autism. Brain.

Lohmann G, Cramon DY von, Colchester ACF. 2008. Deep sulcal landmarks provide an organizing framework for human cortical folding. Cereb Cortex. 18:1415–1420.

Lord S C. Rutter, M. DiLavore, P. Risi, S. Gotham, K. Bishop. 2012. Autism Diagnostic Observation Schedule–2nd edition (ADOS-2). Los Angel CA West Psychol Corp. ed.

Mevel K, Fransson P, Bölte S. 2014. Multimodal brain imaging in autism spectrum disorder and the promise of twin research. Autism.

Oldfield RC. 1971. The assessment and analysis of handedness: the Edinburgh inventory. Neuropsychologia. 9:97–113.

Retico A, Giuliano A, Tancredi R, Cosenza A, Apicella F, Narzisi A, Biagi L, Tosetti M, Muratori F, Calderoni S. 2016. The effect of gender on the neuroanatomy of children with autism spectrum disorders: a support vector machine case-control study. Mol Autism. 7.

Robinson EB, St Pourcain B, Anttila V, Kosmicki JA, Bulik-Sullivan B, Grove J, Maller J, Samocha KE, Sanders SJ, Ripke S, Martin J, Hollegaard MV, Werge T, Hougaard DM, Als TD, Baekvad-Hansen M, Belliveau R, Demontis D, Dumont A, Goldstein J, Grauholm J, Hansen CS, Hansen TF, Howrigan D, Lescai F, Mattheisen M, Moran J, Mors O, Nordentoft M, Norgaard-Pedersen B, Poterba T, Poulsen J, Stevens C, Walters R, Neale BM, Evans DM, Skuse D, Mortensen PB, Børglum AD, Ronald A, Smith GD, Daly MJ. 2016. Genetic risk for autism spectrum disorders and neuropsychiatric variation in the general population. Nat Genet. 48:552–555.

Ronald A, Happé F, Bolton P, Butcher LM, Price TS, Wheelwright S, Baron-Cohen S, Plomin R. 2006. Genetic Heterogeneity Between the Three Components of the Autism Spectrum: A Twin Study. J Am Acad Child Adolesc Psychiatry. 45:691–699.

Rutter M, Lecouteur A, Lord C. 2003. The Autism Diagnostic Interview-Revised (ADI-R). Los Angel CA West Psychol Serv.

Sandin S, Lichtenstein P, Kuja-Halkola R, Larsson H, Hultman CM, Reichenberg A. 2014. The familial risk of autism. JAMA. 311:1770–1777.

Schaer M, Kochalka J, Padmanabhan A, Supekar K, Menon V. 2015. Sex differences in cortical volume and gyrification in autism. Mol Autism. 6:42.

Schumann CM, Bloss CS, Barnes CC, Wideman GM, Carper RA, Akshoomoff N, Pierce K, Hagler D, Schork N, Lord C, Courchesne E. 2010. Longitudinal Magnetic Resonance Imaging Study of Cortical Development through Early Childhood in Autism. J Neurosci. 30:4419–4427.

Shaw P, Kabani NJ, Lerch JP, Eckstrand K, Lenroot R, Gogtay N, Greenstein D, Clasen L, Evans A, Rapoport JL, Giedd JN, Wise SP. 2008. Neurodevelopmental Trajectories of the Human Cerebral Cortex. J Neurosci. 28:3586–3594.

Toro R, Perron M, Pike B, Richer L, Veillette S, Pausova Z, Paus T. 2008. Brain size and folding of the human cerebral cortex. Cereb Cortex. 18:2352–2357.

Van Wijngaarden-Cremers PJM, van Eeten E, Groen WB, Van Deurzen PA, Oosterling IJ, Van der Gaag RJ. 2014. Gender and age differences in the core triad of impairments in autism spectrum disorders: a systematic review and meta-analysis. J Autism Dev Disord. 44:627–635.

Via E, Radua J, Cardoner N, Happé F, Mataix-Cols D. 2011. Meta-analysis of Gray Matter Abnormalities in Autism Spectrum Disorder: Should Asperger Disorder Be Subsumed Under a Broader Umbrella of Autistic Spectrum Disorder? Arch Gen Psychiatry. 68:409.

von dem Hagen EAH, Nummenmaa L, Yu R, Engell AD, Ewbank MP, Calder AJ. 2011. Autism Spectrum Traits in the Typical Population Predict Structure and Function in the Posterior Superior Temporal Sulcus. Cereb Cortex. 21:493–500.

Wechsler D. 2003. Wechsler Intelligence Scale for Children^®^ - Fourth Edition. Pearson.

Wechsler S. D. Coalson, DL. Raiford. 2008. WAIS-IV: Wechsler adult intelligence scale. San Antonio TX Pearson. ed.

Werling DM, Geschwind DH. 2013. Sex differences in autism spectrum disorders. Curr Opin Neurol. 26:146–153.

Zetterqvist J, Sjölander A. 2015. Doubly Robust Estimation with the R Package drgee. 4.

